# Repeated Measures Regression in Laboratory, Clincal and Enviromental Research - Different Between/Within Subject Slopes and Common Misconceptions

**DOI:** 10.1101/374124

**Authors:** Donald R. Hoover, Qiuhu Shi, Kathryn Anastos

**Affiliations:** Department of Statistics and Biostatistics and Institute for Health, Health Care Policy and Aging Research, Rutgers University, Piscataway, NJ, US; School of Health Sciences and Practice/New York Medical College, Valhalla, NY, US; Albert Einstein College of Medicine/Montefiore Medical Center, Bronx

**Keywords:** Between/Within Subject Associations, Repeated Measures, Cross Sectional Regression, Generalized Estimating Equations, Mixed Models, Working Correlation

## Abstract

<u>When using repeated measures linear regression models to make causal inference in laboratory, clinical and environmental research, it is often assumed that the Within Subject association of differences (or changes) in predictor value across replicates is the same as the Between Subject association of differences in those predictor values</u>. But this is often false, for example with body weight as the predictor and blood cholesterol the outcome i) a 10 pound weight increase in the same adult more greatly a higher increase in cholesterol in that adult than does ii) one adult weighing 10 pounds more than a second reflect increased cholesterol levels in the first adult as the weigh difference in i) more closely tracks higher body fat while that in ii) is also influenced by heavier adults being taller. Hence to make causal inferences, different Within and Between subject slopes should be separately modeled. <u>A related misconception commonly made using generalized estimation equations (GEE) and mixed models (MM) on repeated measures (i.e. for fitting Cross Sectional Regression) is that the working correlation structure used only influences variance of model parameter estimates</u>. But only independence working correlation guarantees the modeled parameters have any interpretability. We illustrate this with an example where changing working correlation from independence to equicorrelation qualitatively biases parameters of GEE models and show this happens because Between and Within Subject slopes for the predictor variables differ. We then describe several common mechanisms that cause Within and Between Subject slopes to differ as; change effects, lag/reverse lag and spillover causality, shared within subject measurement bias or confounding, and predictor variable measurement error. The misconceptions noted here should be better publicized in <u>laboratory, clinical and environmental research</u>. Repeated measures analyses should compare Within and Between subject slopes of predictors and when they differ, investigate the reasons this has happened.

**HIGHLIGHTS:** When using repeated measures with time varying predictors variables in laboratory, clinical and environmental research:

1. Cross sectional regressions with any working correlation structure other than independence often give non-meaningful results
2. Between/Within subject decomposition of slopes should be undertaken when making causal inferences
3. Investigators should investigate the reasons Between and Within Subject slopes differ if this occurs

## 1. Introduction

Two common misconceptions made in laboratory, clinical and environmental research fitting repeated measures regression with Generalized Estimating Equations (GEE) and Mixed Models (MM) are: <u>Misconception-A</u>: The association between the predictor variable and outcome across different measures from the same Subject (Within Subject) is the same as the association of that variable with the outcome between measures from different subjects (Between Subject). In fact these associations can differ which should be considered when making causal inference. For example with weight as the predictor and cholesterol the outcome, i) a 10 lb. increase within the same person more likely indicates greater difference in serum cholesterol than does ii) one person being 10 lbs. heavier than another as i) more likely reflects body fat gain while ii) also can indicate that the heavier person is taller. <u>Misconception-B</u>: The working correlation structure used in GEE and MM models is only a nuisance factor that impacts precision of model parameter estimates. As Table 1 in the next paragraph illustrates and Section 2 explains, a wrong choice for working correlation biases parameter estimates. Both of these misconceptions are related to each other, but the analytical details are complicated. So we begin with a direct illustration of <u>Misconception-B</u> that we later show relates to <u>Misconception-A</u>.

Table 1 presents parameter estimates from repeated measures linear regression using a clinical/laboratory measure glomerular filtration rate (EGFR) from the MDRD Formula[1] as the outcome Y and three laboratory predictor variables (X_1_, X_2_, X_3_)=(HIV infection, serum albumin, blood urea nitrogen (BUN)) fit to 10,782 semiannual measures of 584 women at the Bronx-site of the Women’s Interagency HIV Study (WIHS)[2]. Higher EGFR values indicate better renal function. The models fit in this Table assume that the Between and Within Subject associations of the predictor variables are the same; we later show this assumption is incorrect. The parameter estimates of Table 1 were calculated using GEE[3] with both independence (GEE-IND) in columns 2-4 and equicorrelation (GEE-E) columns 5-7 for the working correlation of model residuals of repeated measures in the same person. (The row notation used in this Table for the slope parameters; *β_CS,,HIV_ β_CS,,ALB_* and *β_CS,,BUN_*) is explained later in the paper).

**Table 1 –.**
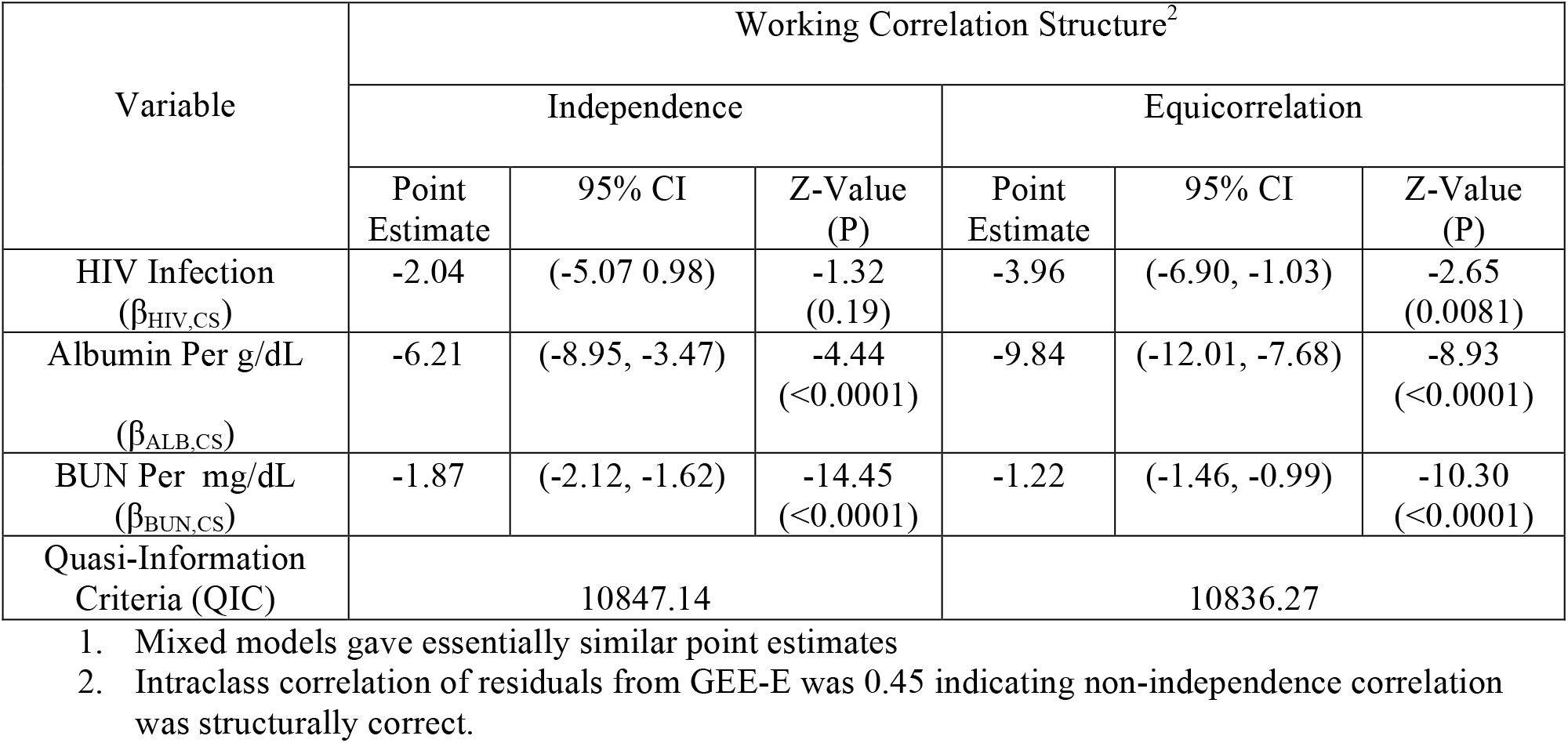
Cross Sectional Regression Using GEE^1^ of EGFR = HIV infection, Serum Albumin and BUN in the Bronx WIHS

| Variable | Working Correlation Structure<sup>2</sup> |  |  |  |  |  |
| --- | --- | --- | --- | --- | --- | --- |
|  | Independence |  |  | Equicorrelation |  |  |
|  | Point Estimate | 95% CI | Z-Value (P) | Point Estimate | 95% CI | Z-Value (P) |
| HIV Infection ($\beta_{HIV,CS}$) | -2.04 | (-5.07 0.98) | -1.32 (0.19) | -3.96 | (-6.90, -1.03) | -2.65 (0.0081) |
| Albumin Per g/dL ($\beta_{ALB,CS}$) | -6.21 | (-8.95, -3.47) | -4.44 (<0.0001) | -9.84 | (-12.01, -7.68) | -8.93 (<0.0001) |
| BUN Per mg/dL ($\beta_{BUN,CS}$) | -1.87 | (-2.12, -1.62) | -14.45 (<0.0001) | -1.22 | (-1.46, -0.99) | -10.30 (<0.0001) |
| Quasi-Information Criteria (QIC) | 10847.14 |  |  | 10836.27 |  |  |

1. Mixed models gave essentially similar point estimates 2. Intraclass correlation of residuals from GEE-E was 0.45 indicating non-independence correlation was structurally correct.

Most of today’s literature providing guidance on fitting repeated measures linear regression (i.e. [4–12]) qualitatively describes working correlation as a “nuisance factor” that does not alter model parameters and states *the working correlation that minimizes variance of parameter estimates should be chosen*. However, in Table 1, the parameter estimate for BUN (per g/dL), from GEE-E, of −1.22; 95% confidence interval (CI) (−1.46, −0.99) is both qualitatively and statistically higher than the corresponding GEE-IND estimate of −1.87; 95% CI (−2.12 −1.62).

For HIV, the parameter estimate of −3.86, p= 0.0081 from GEE-E is qualitatively lower than that from GEE-IND −2.04 and p=0.19. Clearly, changing the working correlation from independence to equicorrelation changes the parameters; thus correlation structure is not merely a nuisance factor. We note that if Mixed Models[3] rather than GEE are used for Table 1, The corresponding parameter point estimates in Table 1 using independence correlation (MM-IND) and equicorrelation (MM-E) are essentially unchanged (data not shown).

Faced with this confusing dilemma, investigators typically go to published guidance on which correlation structure to use. To that end, based on the within subject correlation of residuals being 0.45 in GEE-E (and in MM-E), and the quasilikelihood independence criteria goodness of fit statistic (QIC) = 10,836.27 for GEE-E being smaller than the QIC=10,847.14 for GEE-IND (or the Akaike information criteria goodness of fit statistic (AIC) from a mixed model using equicorrelation (MM-E) of (AIC=94,934.5) being smaller than AIC=99,374.5 from a mixed model using independence (MM-IND) (data not shown)) most articles providing model fitting guidance [4–12] point towards using equicorrelation working correlation.

However, as Section 2 describes in detail, this guidance is problematic; only the parameter estimates obtained by using independence working correlation can have any meaning here as the model did not separately fit Within and Between Subject Slopes. Further as Section 3 explains, if the investigators’ goal is causal inference, then decomposition of the parameters into Within and Between subject associations (or slopes) is needed. Sections 4 and 5 present epidemiological factors that cause Within and Between subject slopes to differ and some consequences of this. The Discussion (Section 6) summarizes and explores further implications for our example and other laboratory, clinical and environmental settings.

## 2. Cross Sectional and Between/Within Subject Linear Models With Repeated Measures

We begin here with some notation. Consider repeated measures on *n* subjects denoted by *i*=*1,2*,…*n*. For most laboratory and clinical analyses the “subjects” will be persons with longitudinal repeated measures. But for environmental analyses “subjects” can be something else such as cities, schools, hospitals, etc. Each subject has *J_i_* different observations enumerated by *j*= 1, …, *J_i_* often taken at times *t_i1_* < *t_i2_* <…. <*t_i,Ji_*, on the same person when person is the “subject”. But for some studies, replicates are taken at the same time, such as from *J_i_* neighborhoods in the same city when city is the “subject”. For *J_i_* constant across *i*, we denote *J*. The observations have continuous outcomes *Y_ij_* and *K* predictor (or exposure) variables *X_ij_* = *X*_1,*ij*_, *X*_2,*ij*_,…*X_K,ij_*. When K=1 we drop the “K” enumeration, using *X_ij_* for the only predictor. Linear models for E[*Y_ij_*| *X_ij_*] or E[*Y_ij_*| *X_ij_*] are fit.

### 2A Cross Sectional Regression (CS) Regression

The most commonly fitted linear model does not separate “Within” and “Between” subject associations and is usually written out as *Y_ij_* = *α* + *β*_1_*X*_1,*ij*_ + *β*_2_*X*_2,*ij*_ +…. + *β*_K_*X_K,ij_* + *ε_ij_*. This is denoted “Cross Sectional Regression” for longitudinal repeated measures and we use this same nomenclature for settings where the repeated measures are not longitudinal. We also add a subscripted “CS” to the β’s to distinguish these slopes from Between subject (BS) and Within subject (WS) slopes defined in Section 2B. The Cross Sectional Regression model here is denoted:

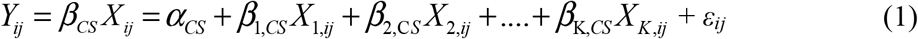

where *α_CS_,β*_1,*CS*_,*β*_2,*CS*_,⋯,*β*_K*CS*_ are unknown parameters, while *ε_ij_* is error with E[ε_ij_]= 0 that is independent between different subjects *i* and *i*’, but may be c–orrelated for *j*≠*j*’ within the same subject. Again for K=1 the subscript for K is dropped; the model is, *Y_ij_* = *α_CS_* + *β_CS_X_ij_* + *ε_ij_*. The goal of Cross Sectional Regression is most appropriately to obtain estimates 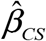 for *β_CS_* to input 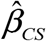 into (1) order to estimate unobserved Y’s from observed *X_ij_*’s as 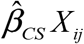. Cross Sectional Regression is also used to make adjusted (causal) inference on the covariate associations in 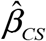 but as we show later, doing this may be problematic.

### 2B Between/Within Subject Slope (BS/WS) Regression

While regression models fit in laboratory, clinical and environmental studies typically do not consider this, it has long been noted that slopes on changes of *X_ij_* within the same subject *i* differ from cross sectional associations[13–16]. To illustrate this consider the cross sectional model of a laboratory measure cholesterol (*Y_ij_*) on the clinical outcome of body weight (*X_ij_*); *E*[*Y_ij_*] = *α_CS_* + *β_CS_ X_ij_*. As described in the first paragraph of the Introduction, the cross sectional slope *β_CS_* for association of a 10 lb. weight difference between two different subjects (i.e. persons) on cholesterol is less than the slope for association of a 10 lb “Within Subject” weight change for the same person on cholesterol which we denote as *β_WS_*. Again the reason *β_CS_* is less than *β_WS_* is; a) part a 10 lb. cross sectional weight difference between two subjects often reflects greater height in one of the persons b) but a 10 pound weight increase in the same subject is not influenced by height difference and thus the is more likely due to more body fat being in the heavier weight. Thus since greater body fat is what is directly associated with more cholesterol, the within person association of a 10 lb. increase with cholesterol is greater than the cross sectional repeated measures association of a 10 lb. difference. Common within person body height reflects a shared *within subject measurement bias* on weight as a predictor of cholesterol. For example as Figure 1a illustrates, if *TX_ij_* = body mass index (*wt*/*ht*^2^) were the true predictor of *Y_ij_* and *H_i_* = height (which does not change with *j* in the same *i*), then *X_ij_* = *TX_ij_* * *H_i_* contains this shared within subject measurement bias *E_i_*. Section 4 describes more settings where *β_WS_* ≠ *β*_CS_.

**Figure 1.**
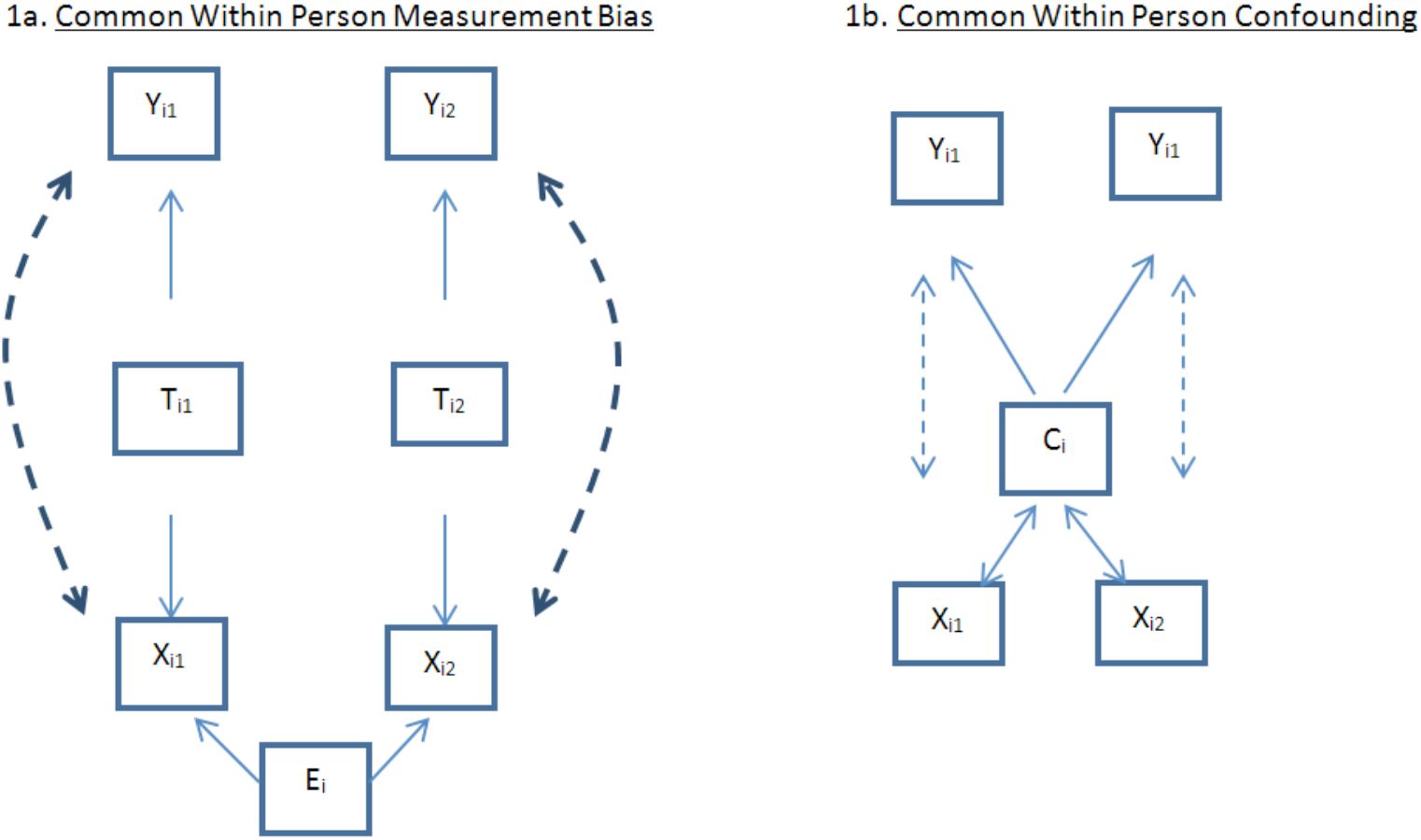
Illustration of Common Within Person Measurement Bias and Confounding for K=1

Therefore linear regression models fit to make causal inference often decompose the associations into “Within Subject” slopes (*β_WS_*) described above and “Between Subject” slopes (*β_BS_*) described below that capture associations of subjects’ central tendencies of the exposure. To do this, subject means of the predictor variables 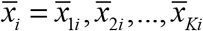 are calculated, where 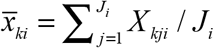. Then *Y_ij_* is modeled as a combination of “Between Subject” slopes from 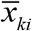 and “Within Subject” slopes from deviations of *X_kij_* about 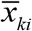.

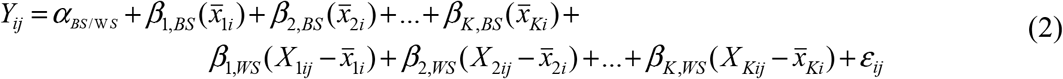

Or for K = 1,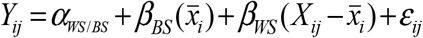. To illustrate this for our earlier example with Y_ij_=Cholesterol and X_ij_=Weight, let *α_WS/BS_* = 30, *β_BS_*=0.9 and *β_WS_*=3; then 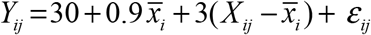. If person *i* had an average value of 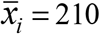 across all *J_i_* measures with the *j^th^* measure being *X_ij_*=200 then for that person-visit *t_ij_*, *E*[*Y_ij_*] =30+0.9(210)+3(200 − 210) =189.

Now for some technical asides; First – the choice of the observed 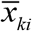 as the “central tendency” of *X_kij_* for subject *i* is necessary as “μ_ki_” a person’s “true average weight” over the entire time period is unknown, but for *J_i_* large enough, 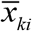 should be close to μ_ki_. Thus while *β*_k,*WS*_ only captures association with within subject change in in *X_kij_, β*_k,B*S*_ inherently contains some *β_k,WS_* from deviation of 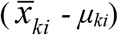; especially for small *J_i_*. Second - the implicit assumption that *β_k,WS_* is well defined may also not always be true. For example “*β_k,WS_*” could differ by time separation t_j_ - t_j_. Perhaps weight gain of 10 lbs. in one month creates a shock that hyper-elevates cholesterol, but a 10 lb. weight gain over 12 months does not; in which case *β_WS_* | *t_j_* − *t_j′_* = 1 > *β_WS_* | *t_j_* − *t_j′_* = 12. But it is probably reasonable to assume that any such differences are minor.

In spite of these technical limitations, Between/Within-Subject decomposition is used including to test if *β*_kB*S*_ = *β*_k,*WS*_ in which case as shown in Section 2C they also equal *β*_k,*CS*_ and the separated *BS, WS* decomposition can be collapsed. Due to orthogonal decomposition of *X_kij_* about 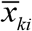, this previous test for collapsing the *BS, WS* decomposition is a two sample z-test of parameter estimates from fitted models comparing 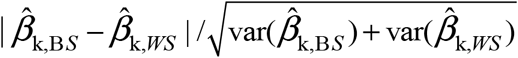 to *Z_1−α/2_* [16]. Between/Within subject decomposition is mostly used for inference on adjusted (causal) associations of the *X_k,ij_*’s on *Y_ij_*’s. It is typically not used to produce models to estimate future unknown *Y_ij_* from as such is often done when only one observation per subject is available; hence 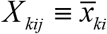.

We refit the analyses of Table 1 to illustrate that the impact of choice of correlation structure (i.e GEE-IND Vs. GEE-E working correlations) is eliminated after making a Between – Within Subject decomposition. There were no new HIV infections after study entry; so 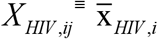 making Within Subject association of HIV unmeasurable. For Within Subject associations of BUN and Albumin, GEE-IND and GEE-E gave identical point estimates as centering about 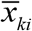 makes comparisons entirely within subject and invariant to these correlation choices (although within subject estimates would differ slightly if autoregressive (AR(1)) or other formulations for intrasubject correlation of residuals had been used). There were small GEE-IND, GEE-E differences on the Between Subject slopes as has been observed elsewhere[17]. For example, the point estimate for between subject HIV status is −1.16; 95%CI (−4.21, 1.88) in the GEE-IND of Table 2 Vs. −1.57; 95% CI (−4.47, 1.33) with GEE-E.

**Table 2.** Between Subject and Within Subject Decomposition Regression Using GEE^1^ for EGFR = HIV Infection, Albumin and BUN in the Bronx WIHS

| Variable | Compartment | Independence Working Correlation |  |  | Equicorrelation Working Correlation |  |  |
| --- | --- | --- | --- | --- | --- | --- | --- |
|  |  | Point Estimate | 95% CI | Z-Value (P) | Point Estimate | 95% CI | Z-Value (P) |
| HIV Infection | Between Subject ($\beta_{HIV,BS}$) | -1.16 | (-4.21, 1.88) | -0.75 (0.45) | -1.57 | (-4.47, 1.33) | -1.06 (0.29) |
|  | NA<sup>2</sup> | --- | --- | --- | NA<sup>2</sup> | --- | --- |
| Albumin Per g/dL | Between Subject ($\beta_{ALB,BS}$) | -3.27 | (-7.88, 1.33) | -1.39 (0.16) | -2.71 | (-7.00, 1.57) | -1.24 (0.21) |
| | Within Subject ($\beta_{ALB,WS}$) | -10.70 | (-12.99, -8.40) | -9.16 (<0.0001) | -10.70 | (-12.99, -8.40) | -9.16 (<0.0001) |
| BUN Per mg/dL | Between Subject ($\beta_{BUN,BS}$) | -2.72 | (-3.10, -2.33) | -13.89 (<0.0001) | -2.65 | (-3.01, -2.08) | -14.21 (<0.0001) |
| | Within Subject ($\beta_{BUN,WS}$) | -1.11 | (-1.34, -0.88) | -9.31 (<0.0001) | -1.11 | (-1.34, -0.88) | -9.31 (<0.0001) |

1. Mixed models gave essentially similar point estimates 2. There is no Within Subject Variation for HIV Infection Status

From now on, we only examine GEE-IND results for Between/Within subject decomposition models as GEE-E results are essentially similar (data not shown). For BUN, the GEE-IND the Between Subject 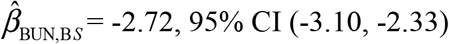 is qualitatively and statistically farther from 0 than the corresponding Within Subject slope; 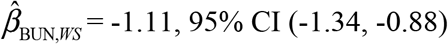. But serum albumin goes the other way; the Within subject slope 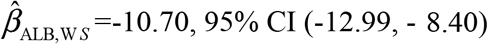 is statistically further from 0 than the corresponding Between subject GEE-IND 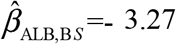 with a 95% CI (−7.88, 1.33) that overlaps 0.

So how can one interpret differences in the Between and Within Subject slopes in particular for causal inference? This depends on hypotheses of interest (and we do not have any for this illustrative example). But general rules also apply although we are unaware of any systematic exploration of reasons that between *β*_B *S*_ (or *β*_B *S*_ for *K*=1) could differ from within subject *β*_W *S*_ (or *β_WS_*) slopes and implications for causal inference. But before going to this here it is important to first note the relationship between, Cross Sectional, Within Subject and Between Subject slopes.

### 2C Relationship Between *β_CS_, β_WS_* and *β_BS_*

*β_CS_* averages *β_WS_* and *β_BS_* according to relative variances of the within subject means Vs. the variance of the repeated measures about those sample means[16]. For example, with *K*=1, if 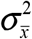 is the population variance of the within person mean 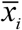 and 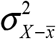 is the population variance of the deviations of differences of the repeat measures *X_ij_* from their 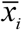, then

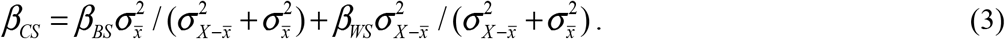

In the previous example of weight and cholesterol with 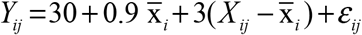, if 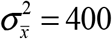 and 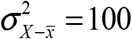, then *β_CS_* =0.9*400/(100 + 400) + 3*100/(100 + 400) = 1.32. If the sample means are more homogeneous in weight with 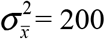 and 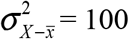 still, then *β_CS_* moves closer to *β_WS_*; *β_CS_* =0.9*200/(100 + 200) + 3*100/(100 + 200) = 1.60.

### 2D Working Correlation Structures for Model Residuals Other than Independence Can Lead to Unusable Results for Cross Sectional Regression

As noted earlier, fitting both MM and GEE repeated measure regression models involve specification of correlation (or working correlation) of *ε_ij_* within the same subject *i*. We denote the working correlation structure by ***V_i_*** which is a matrix. Typical choices for ***V_i_*** are equicorrelation (E) with correlation of *ε_ij_* and *ε_ij’_* for j ≠ j’ always the same value ρ and independence (IND); with correlation of ε_ij_ and ε_ij_’ ≡ 0 as used in the illustrative examples of Table 1 and Table 2, and also AR(1) where correlation of *ε_ij_* and *ε_ij_*’ is *ρ*^|*j*−*j*’|^ [3]. Again, current guidance articles[4–12] emphasize choosing the ***V_i_*** that most closely fits the true covariance structure of the residuals within *i* and/or by model fit criteria such as having lowest QIC for GEE and AIC for MM, as doing so often improves precision of the model parameter estimates. However, this approach may be wrong for Cross Sectional Regression, because using any correlation structure other than independence can introduce structural bias into 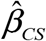 that the standard approaches used to chose the best working correlation structure such as intrasubject residual correlations, the AIC and QIC mentioned earlier do not account for[18, 19].

From Pepe & Anderson (1994)[18], for Cross Sectional Regression if *X_ii_*.varies within *i* and,

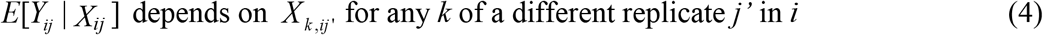

then *no matter what true correlation structure of ε_ij_ among repeated measures within a subject is*, GEE-IND gives unbiased estimates for *β_CS_*, but any MM or GEE model not using ***V_i_*** = Independence, gives biased estimates of *β_CS_*. Thus the only working correlation structure that should be used to estimate *β_CS_* is ***V_i_*** = IND. However, if (4) does not hold, then any working correlation structure obtains unbiased estimates for *β_CS_* in which case, choosing the ***V_i_*** that most accurately gives the correlation structure of *ε_ij_* minimizes the variance of 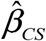.

If repeated measures *j* and *j*’ are thought of as “siblings” and the predictors as “exposures” then (4) means that even after considering the “self-exposure” of the current measure *j* through *X_ij_*, the outcome *Y* has “**<u>Co</u>**nditional **<u>D</u>**ependence **<u>O</u>**n **<u>S</u>**ibling **<u>E</u>**xposures” (i.e. on *X_k,ij’_*). Thus the sibling exposure *X_k,ij’_*, could be thought of as a “Co-DOSE” beyond the “dose” from the “self-exposure”. Hence we use “Co-DOSE” to denote that (4) occurs.

Also while this point has not very well made, for Cross Sectional Regression, Co-DOSE or (4) largely occurs if and only if Between and Within Subject slopes differ. If Between and Within Subject slopes differ for any predictor (i.e. *β_BS_* ≠ *β_WS_*) then Co-DOSE (4) happens. But if the Between and Within Subject slopes are equal for all predictors (i.e. *β_BS_* = *β_WS_*) then (4) does not occur. More details on this and an illustration are given in Appendix 1. But one trivial case is if the predictors are invariant within the same subject; (i.e. 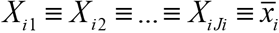 then the Within Subject slopes are not defined since 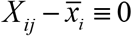and for the same reason Co-DOSE cannot occur. While the mathematical details are beyond this paper, if *β_BS_* ≠ *β_WS_*, when **V_i_** ≠ *IND*, the non-zero covariances *ρ_ij_* > 0 besides adjusting for within *i* collinearity of *ε_ij_ also* overweight the *β_BS_* relative to *β_WS_* in (3) pushing “Cross Sectional Regression” parameter estimates away from *β_CS_* towards *β_WS_* [16]. Since robust covariance methods exist to adjust for impact of misspecification of ***V_i_***=IND from collinearity of the *ε_ij_* on variance estimates in particular for GEE[3], **V_i_** = *IND* can eliminate bias in estimating *β_CS_* and still provide a conservative variance for the estimate.

### 2E Implications for Clinical, Laboratory and Environmental Research

Much of what has been presented above is not commonly understood and implemented in clinical, laboratory and environmental research. Cross sectional models are typically fit, with *β_CS_* interpreted to also be *β_BS_* and *β_WS_* without checking if these slopes are equal. Non-independence ***V_i_*** is often used for Cross Sectional Regression without checking if Co-DOSE (4) exists. Systematic epidemiological description of causal mechanisms for why Between and Within subject slopes can differ for clinical, laboratory and environmental measures have not been undertaken. We thus do this next in Section 3.

## 3. Classification of Epidemiological Reasons for Between and Within Subject Slopes to Differ

To make it easier for investigators to identify what could cause *β_k,WS_* ≠ *β_k,BS_* (or equivalently Co-DOSE) in given settings, (especially for clinical, laboratory and environmental) we classify major reasons this happens. For simplicity now let *K*=1 unless otherwise noted as the principles below extend to multivariate settings.

### 3A. Change Effects

We propose that the effect of a longitudinal within subject change in the predictor X could have a greater (or less) direct impact on Y than a long term standing difference in X between two different subjects (hence *β_WS_* ≠ *β_BS_*) and define this as a (c.f. short term) “Change Effect”. Returning to the example of weight and cholesterol, consider two identical twins, A has lived his adult life at 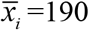 lbs. and B at 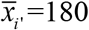 lbs. If B undergoes a short term weight gain of 10 lbs. to 190 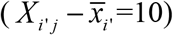, assuming 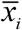 not impacted by the rapid change, while A remains at 190 lbs. 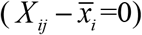, the shock or corollaries of this rapid change in “B” may raise his cholesterol level above that of “A’s” even though they both now weigh 190 lbs. meaning *β_WS_* > *β_BS_* and Co-DOSE (4) occurs. But it should be noted that as was mentioned in Section 2B, in this setting, *β_WS_* would be undefined if say a 10 lb. gain in a shorter time period (i.e. 1 month) increases *β_WS_* more than does a 10 lb. gain over a longer time period (i.e. 12 months).

### 3B Lag Causality of *X* on Future *Y*

The effect of prior levels of *X* on *Y* may independently project into the future (i.e. beyond that effect of the current level of X). For example, consider an HIV infected person and two time points *t_1_* < *t_2_*; let *X_ij_* be HIV viral load and *Y_ij_* be CD4 count. High HIV levels destroy CD4 blood cells into the future. So as illustrated in Figure 2a, high HIV viral load at *t_1_* may affect CD4 loss from t_1_ to t_2_ (*lag causality of X at t_1_ on Y at t_2_*) so that even if the person’s HIV viral load is low at t_2_, the high viral load at t_1_ is predictive of lower CD4 at t_2_ through that higher viral load at t_1_ having created more CD4 destruction between t_1_ and t_2_. Thus *Y_i,2_*|*X_i2_* at *t_2_* is not independent of *X_i,1_* at t_1_; Co-DOSE (4) occurs and Within / Between Subject slopes differ; *β_BS_* ≠ *β_WS_*. Lag causality is often considered when serial measures of *X* = long term environmental exposures (such as air pollution) that effect chronic conditions *Y* (such as lung function) are obtained[3,20].

**Figure 2.**
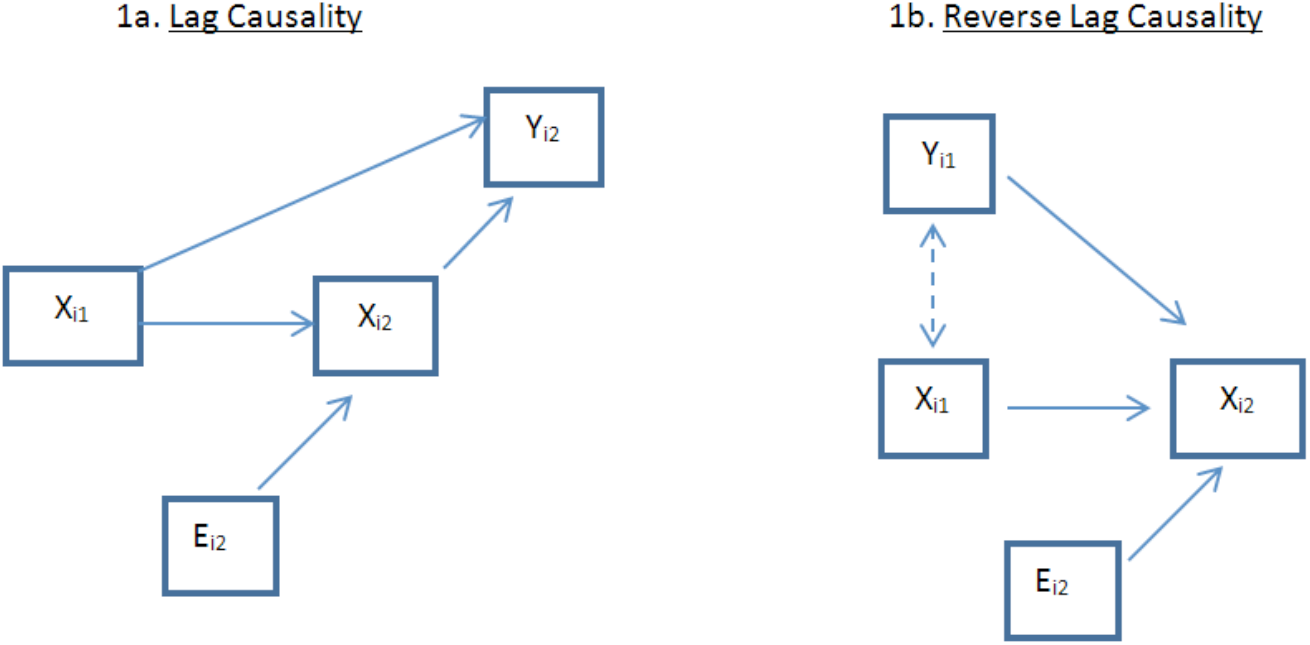
Illustration of Lag Causality and Reverse Lag Causality for K=1

### 3C Reverse Lag Causality of *X* on Future *Y*

The setting in Section 3B also manifests in the opposite direction if *X* is being used as to estimate *Y* that is causal for future *X*. Reversing the previous example with *X* now being CD4 used to predict HIV viral load as Y, as Figure 2b illustrates, high viral load (*Y_i,1_*) at t_1_ may have degraded the CD4 count from t_1_ to t_2_. Thus *Y_i,1_*|*X_i1_* at *t_1_* is not independent of *X_i,2_* at *t_2_*; Co-DOSE (4) occurs and Between / Within subject slopes differ; *β_BS_* ≠ *β_WS_*.

### 3D Spillover Causality of X on Adjacent Y

The setting of 3B can also manifest in repeated measure cross sectional settings based on geographical proximities. Let the subjects (*i*) now be cities and j enumerate different neighborhoods in these cities. The repeat measures are *X_ij_* = average air pollution of neighborhood *j* in city *i* and *Y_jj_* = average lung function of all residents living within neighborhood *j* of city *i*. A resident living in neighborhood *j* may work in different neighborhoods *j*’ of the same city and thus have “*spillover exposure*” to air in the neighborhood they work in, for a given city i, *Y_i,i_*|*X_ij_* is not independent of *X_ij_*’ and hence Co-DOSE occurs.

### 3E Common within Subject Measurement Bias

Shared *within subject measurement bias* occurs if all repeat measures from the same subject have the same positively correlated measurement bias. This was the setting described in Section 2B and Figure 1a with weight as exposure for cholesterol. Here with weight as a surrogate for body fat, the measurement bias was mediated by height with taller people being heavier independently of body fat than shorter persons which lead to *β_WS_* > *β_CS_* and Co-DOSE (4) when weight was a predictor of cholesterol. In this setting height is a *measurement bias* not as a confounder as height itself is not associated with cholesterol. We now present a similar setting where the unmeasured variable is a confounder.

### 3F Common within Subject Confounding

Figure 1b shows a diagrammatically similar phenomenon, that causes *β_WS_* ≠ *β_CS_* and Co-DOSE (4), *common within subject confounding* where now the extraneous factor shared by the repeated measures of the same subject is associated with Y. For example, let the confounder variable *C_i_* = sex of subject *i* (which does not change with j) not be in the model and the outcome *Y_i,j_* be a linear score for male pattern baldness at time j with again *X_ij_* being weight at time j. Now men are both heavier and independently of weight have greater male pattern baldness than women. So *C_i_* is associated with the outcome. Here a 10 lb. weight difference in 2 persons, but not a within person increase of 10 lbs., could be informative of the heavier person more likely being male. Hence for this example, *β_W_S* = 0 (assuming within person weight does not influence baldness), but *β_BS_* > 0 and hence also *β_CS_* > 0 reflecting unaddressed between subject confounding from heavier persons more likely to be men. Similarly, Mancl, Leroux & DeRouen proposed that unmeasured treatment compliance as a confounder could lead to Between/Within subject slope differences when *j* enumerated different teeth in the same subject[17].

In non-longitudinal settings where *i* denotes clusters (for example schools) and *j* denotes repeated subjects within that cluster (for example students), common within subject confounding is referred to as “contextual effects”[21,22]. For example as Robinson (1950)[13] observed, when X was race of the student (White=0, Black=1) and *Y* was achievement-score, a higher 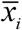 (portion of a school’s students that were non-White) indicated weaker financial support for that school (weaker financial support being the confounder) and thus worse achievement-scores overall for that school; *β_BS_* was negative. But within the same school Black and White students performed equally well, thus *β_WS_* was 0. Begg & Parides (2003) identify a similar setting in birthweight and IQ in families[23].

### 3G Measure Error in *X_ij_* Makes E[*Y_ij_*|*X_ij_*] Dependent on *X_ij’_*

It has been shown that, measure error on *X* that is independent of *Y*[24] or correlated with Y[25] biases estimates for association of X with Y. Now measure error can arise either from *imprecision in an analysis instrument*, such as in a machine quantifying components of serum, or *in data collection process*, such as the chemical composition of blood samples being non-informatively influenced by diurnal and other nuisance processes. If *X_ij_* is incorrectly quantified due to such measure error, then Co-DOSE (4) occurs and Within / Between subject slopes differ, *β_BS_* ≠ *β_WS_*, because the biases being created from the measure error distribute differentially to different slopes. Figure 3 shows, if *X_i1_* incompletely measures the true state *TX_i1_* (i.e. true BUN) then *X_i2_*, is informative for *TX_i1_* even after considering *X_i1_*.

**Figure 3.**
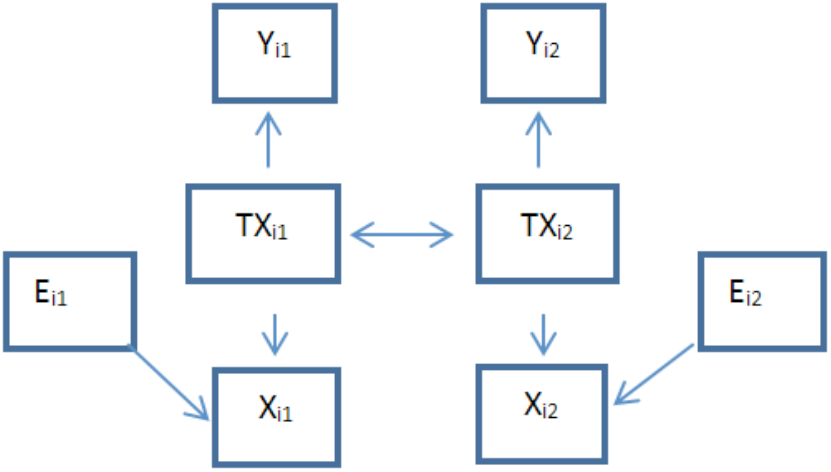
Illustration of Residual Association With Independent Measure Error in X for K=1

For example, Going back to the analysis of Table 1, let *X_ij_* = BUN and *Y_ij_* = EGFR. Let two persons have BUN of *X_i1_*=10 mg/dL measured today but with Measure Error in BUN Assume the true BUN state changes slowly and after 6 months one of these persons measures *X_i2_*=20 mg/dL while the other measures *X_i2_*=5 mg/dL. Since BUN changes slowly, it is more likely that the True BUN today *(TX_i1_)* of the former person is > 10 mg/dL and of the later is < 10 mg/dL. Thus, since i) EGFR (*Y_i1_*) directly depends on *TX_i1_* not *X_i1_*, and ii) *X_i2_* is informative on *TX_i1_* after considering *X_i1_*, thus iii) *Y_i1_*|*X_i1_* is not independent of *X_i2_* and similarly *Y_i2_*|*X_i2_* not independent of *X_i1_* meaning Co-DOSE (4) occurs and Between/Within subject slopes differ. As Appendix 2 shows, measure error on the exposure that is independent of the outcome pushes both *β_WS_* and *β_BS_* towards 0, but more so for β_WS_. Such tempering from averaged measure error has been proposed as a reason |*β_WS_*|<|*β_BS_*| was observed in dental research[17].

But if *M_ij_* is correlated with *Y_ij_* (most likely correlated with measure error on *Y_ij_*[25]) the tempering of *β*’s from *M_ij_* will not be to 0. For example, consider *TX*=CD8 and *TY*=CD4 cells which together are the almost exclusive components of serum lymphocytes (*TZ*)…. (i.e., *TY* ≈ *TZ − TX*). Physiologically *TZ* is constrained creating negative *β_BS_, β_WS_* and *β_CS_* for *TY_ij_* on *TX_ij_*; subjects with higher CD8 component of serum lymphocytes by converse having lower CD4 components. But the measured lymphocyte count (*Z*) is subject to a correlated measure error that equally spreads onto *X* and *Y*; for example if a person is dehydrated, the entire lymphocyte (meaning both CD8=X and CD4=Y) portions of blood become higher. If a person-sample has a high (or low) measured lymphocyte count *Z_ij_* = T*Z_ij_*+*M_ij_*, due to such sampling/measurement error, *M_ij_* distributes onto both CD4 (*X_ij_*) and CD8 (*Y_ij_*) components making both simultaneously higher (or lower). Thus within person, higher measured CD4 count due to positive *M*_*ij*_ is associated with higher measured CD8 as the “*M*_*i*j_” is shared making *β_WS_* drawn towards being positive. But *β_BS_* that tempers down *M_ij_* on both *X* and *Y* through averaging as shown in Appendix 2 for *X* is less affected.

## 4. Predictors having Co-DOSE will bias adjusted parameter estimates of predictors not having Co-DOSE in cross sectional regression when *V_i_* ≠ Independence is used

Going back to Table 1, it was shown earlier that the adjusted point estimate from GEE-IND 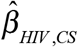 for the adjusted cross sectional association of HIV with EGFR is still consistent for *β*_HIV,*CS*_. However, HIV infection status was constant over all replicates within the same subject so cannot have Co-DOSE (4) as the entire effect of HIV is mediated between subject, not within subject so the question arises whether the adjusted estimate from an non-independence correlation structure 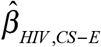 can be biased for *β*_HIV,*CS*_. Note, that for this section, we use 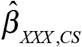 and 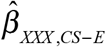 to denote estimates for adjusted cross sectional association for variable XXX from models using independence and equicorrelation structures, respectively. <u>The added designation of “E” (CS-E) in the subscript for equicorrelation but none for independence correlation is made because the equicorrelation estimate (but not the independence estimate) can be biased</u>. The specific question addressed here is could having BUN and albumin that each have Co-DOSE (4) in the model bias the corresponding estimate for cross sectional adjusted HIV association from using equicorrelation 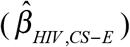 so that it no longer is consistent for β_HIV,CS_ in the multivariate model even though HIV itself is not Co-DOSE? This is important as 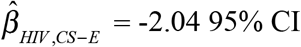 (−5.07 0.98) qualitatively differs from 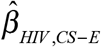 (−6.90, −1.03) in Table 1 with only 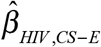 statistically (P < 0.01) differing from 0.

We believe 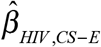 for HIV is biased away from *β_HIV,CS_* and to make this point refer to Table 3 which presents normative data broken down by HIV status of the subjects. A) From Table 1 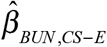 is biased higher for *β_BUN,CS_* (with GEE-E 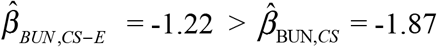 for GEE-IND), while from Table 3, those who are HIV+ have higher mean BUN (12.94 Vs. 12.10, p <0.0001 from GEE-IND). Thus the full “negative effect” of the higher BUN in HIV+ subjects from *β_BUN,CS_* is underestimated by 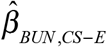 and this pushes 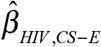 down to compensate. B) Similarly, from Table 1, 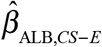 is biased lower for *β_ALB,CS_* (with GEE-E 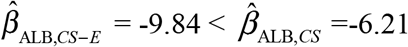) while from Table 3, HIV+ individuals have lower mean albumin; (3.97 Vs. 4.10, P < 0.0001 from GEE-IND). Thus the “positive effect” of the lower albumin in HIV+ subjects from *β*_ALB *CS*_ is overestimated by 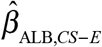 which pushes 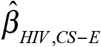 further down to compensate. C) Thus as Figure 4 shows, these deficits in A) and B) above are added to push 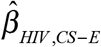 downwards from the true adjusted *β_HIV,CS_*. Therefore, non-independence ***V_i_*** can bias multivariate cross sectional parameter estimates of variables that do not carry Co-DOSE (4) if other variables in the model carry Co-DOSE.

**Figure 4.**
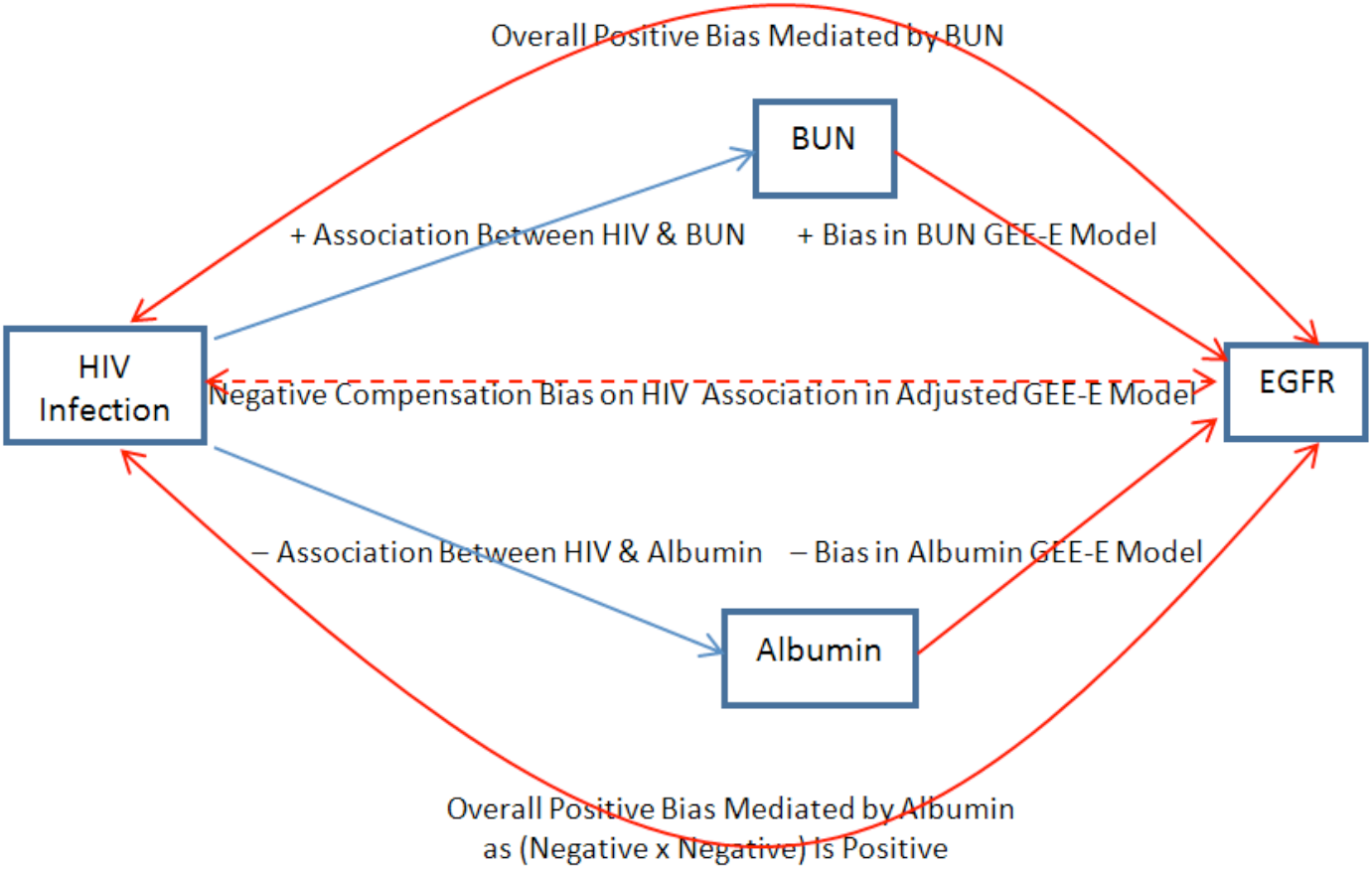
Compensating Bias Pathways on Time Invariant FI IV Estimate From Failure to Use An Independent Working Correlation Structure in Repeated Measures GEE

**Table 3.** Means ± Standard Deviation of EGFR Serum Albumin and BUN Broken Down by HIV Status Across all Repeated Measures Used in Tables 1 and 2

| Variable | For HIV+ Subjects (496 persons<br>7,326 Replicates) | For HIV- Subjects (178 persons<br>3,456 Replicates) |
| --- | --- | --- |
| EGFR | 90.3±27.2 | 92.4±25.0 |
| BUN | 12.94±5.71 | 12.10±5.30 |
| Albumin | 3.97±0.44 | 4.14±0.36 |

## 5. Discussion

New GEE and MM cross sectional regression models from repeated measures of predictors that vary within subject that either use non-independence working correlations or do not state the correlation structure used continue to be published in the clinical, laboratory and environmental sciences literature. These papers do not show awareness of the points presented in Sections 1–4. Specifically, they either; a) do not specify if the coefficients of interest are *β_CS_*, *β_WS_* or *β_BS_* nor check if *β_BS_* = *β_WS_*, b) make potentially invalid *β_CS_* interpretations from MM and GEE using non-independence correlation ***V_i_***’s, and/or c) do not justify the choice of nonindependence working correlations *V_i_* in light of potential differences between *β_WS_, β_BS_* and *β_CS_*. We found >30 such papers from a quick literature search (including some authored by us before becoming aware of these issues); this is probably only a fraction of the total.

Yet papers published up to 65 years ago either warn against using non-independence working correlation structure in cross sectional regression with repeated measures[17,18], or to decompose the associations into Within Subject (*β_WS_*) and “Between Subject” (*β_BS_*) to make causal inference[13–16]. Numerous examples where *β_CS_* ≠ *β_BS_* ≠ *β_WS_* have been presented[13–18,20–23]. While it was not covered in our paper, this includes fitting GEE models of binary outcomes where the issues discussed here also apply[17,26]. But these points are still not well known or emphasized in statistical software documentation and papers providing guidance on GEE and MM analyses (i.e.[4–12]).

One problem that impedes acceptance of Within Subject (*β_WS_*) and “Between Subject” (*β_BS_*) decomposition is that it leads to much more complicated models that are very difficult to explain. Still some studies in environmental research have considered one mechanism described in Section 3. For example, lag causality has been considered in longitudinal analyses of association of air pollution on health measures[3]. Other air pollution / health studies have fit Within and Between Subject decompositions using cities as the subject and neighborhoods as the repeated measures within the city[27–30]. Most often in these studies the magnitude was greater for Within subject slope | *β_WS_* | > | *β_BS_* |, but sometimes | *β_BS_* | > | *β_WS_* | meaning possibly multiple etiologies are involved. Those papers that did attempt to explain the reasons for the differences described common within subject confounding (Section 3E), such as unmodeled pollutants that were correlated between (but not within) cities with the modeled pollutants of interest as a potential reason.

We concur with others[17,18], that Cross Sectional Regression with repeated measures should use independence as the default working correlation unless justification is given to use other ***V_i_***. While non-independence ***V_i_*** can improve precision [20] they can considerably bias estimates for cross sectional parameters *β_CS_*; including perhaps towards what the investigator wants to see. For example in Table 1, P < 0.01 for was observed association with HIV in GEE-E compared to the more appropriate P=0.19 from GEE-IND.

While showing this is beyond the scope of our paper, when ***V_i_*** is not independence, factors such as *J_i_* and magnitude/structure of *ε_ij_* strongly influence parameter estimates for *β_CS_* from the misfitted Cross Sectional Models allowing the misfitted estimate to arbitrarily range from *β_WS_* to *β_CS_*. Standardization is important; and as such factors will arbitrarily vary between studies, parameter estimates of *β_CS_* become harder to compare across studies when ***V_i_*** differs at discretion of investigators. The working correlation structure used in Cross Sectional Regressions using repeated measures should thus always be reported.

We also concur with others [13–17,21–23] that in spite of the difficulties in identifying why “Within” and “Between subject slopes differ, causal inference analyses with repeated measures should initially make such decompositions and investigators be wary if there are qualitative differences between *β_BS_* and *β_WS_*. The example, Table 2 with 584 subjects and 10,782 measurements demonstrated need for *β_BS_, β_WS_* decomposition to make causal inference (as well as for using GEE-IND in cross sectional regression). But a smaller study could have been less clear-cut. If the same point estimates for *β_BS_* and *β_WS_* seen in Table 2 were observed but did not statistically differ, one would be tempted to merge *β_BS_* and *β_WS_* into a combined *β_CS_* at least for some variables as standard model fitting practice promotes parsimony when statistical significance is not observed. This would be particularly true if for a given variable, *k*, neither 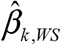 nor 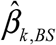 statistically differed from 0, but 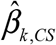 did. If such collapsing is done, it may still be important to report 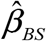 and 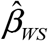 for comparison to future studies and target potential mechanisms for Between/Within slope differences as described in Section 3.

Unfortunately, Between/Within Subject decomposition expands required analyses and presentation. Statistical software mostly does not have standard subroutines to do this. Decomposition can be tedious if 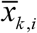 is recalculated to maintain orthogonal decomposition of *X_k,ij_* as new models are fit if observations are excluded from the *J_i_* due to missing values of newly included variables. The fact that 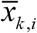 is ill-defined by averaging the *X_k,ij_* rather than being a true mean for subject *i* creates confusion about interpretation of 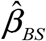 which can also be influenced by within subject slopes as noted in Section 2B.

When 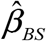 and 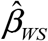 differ, the causal mechanisms as to why this happens should be explored. For example, in Table 2 with EGFR as the outcome, for BUN the Between Subject slope 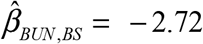 (from GEE-IND) was statistically farther from 0 in the expected direction of association than was the Within Subject slope 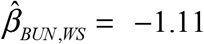. But albumin went the other way; Within Subject slope 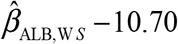 was statistically farther from 0 than was Between Subject slope 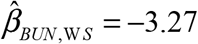. So what are the potential reasons for this?

While lag/reverse lag causality between serum BUN and creatinine (the main component of calculated EGFR) which could reduce magnitude of *β_BUN,WS_* Vs. *β_BUN,BS_*(Sections 3B and 3C), this was unlikely given separation of visits was 6 months and internal biochemistry operates over shorter time periods. But independent measure error on BUN (Section 3E) would temper |*β_BUN,WS_*| relative to |*β_BUN,BS_*|. Several articles find greater; coefficient of variation [31,32], within person change[31,32], assay error[32], and sample degradation for serum BUN Vs. albumin measures[33], all of which could reflect BUN having larger independent measure error than does albumin that would selectively temper 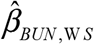 (i.e., more than 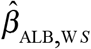) to 0.

Conversely, serum creatinine and albumin are both constrained into the intravascular fluid compartment and will noninformatively increase together with greater hydration and decrease with less hydration of this compartment inducing positively correlated measurement error; much as was the case for measured CD4 and CD8 cells in the last paragraph of Section 3F. As creatinine factors inversely into EGFR calculation, this would constitute negative correlation of measure error between albumin and EGFR and selectively bias 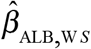 to be more negative than 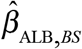. However, BUN, which crosses across all body compartments, is less subject to such correlation in measure error with creatinine and thus with EGFR.

When Between and Within subject slopes differ, (*β_BS_* ≠ *β_WS_*) it is unclear which is the “least confounded/biased”, including the possibility that by “averaging” the different biases in each that *β_CS_* could be least biased. There may be a heuristic perception to believe that by “matching within the same subject”, *β_WS_* is superior to *β_BS_* and *β_CS_*, but this is not necessarily true as Measure Error on *X* (Section 3F) and lag / reverse lag and spillover causality (Sections 3B–3D) can in fact bias *β_WS_* to a larger degree than happens for *β_BS_* and *β*_CS_.

To conclude, for decades it has been known by some that when exposures vary within subjects in repeated measures regression then, i) Cross Sectional Regression using ***V_i_***=Independence working correlation should be the default if building a model to estimate a future unknown *Y* is the goal, and ii) Between/Within subject decompositions of slopes should at least initially be fit when building models for causal inference. Yet this advice rarely makes it into published guidelines and hence is not heeded; perhaps in part due to complexity of the settings where Between and Within Subject slopes differ and limited substantive study of the mechanisms that cause such differences. Clinical, laboratory and environmental studies should explore and quantify reasons for bias that occur in order to provide groundwork to improve future studies. To that end, analyses using repeated measures regression should investigate if Between and Within Subject slopes differ and when they do, try to identify the reasons for this including; change effect, lag/reverse lag and spillover causality, shared within subject measurement bias or confounding, and/or measure error on exposures.

## Acknowledgements

Data in this manuscript were collected by the Women’s Interagency HIV Study (WIHS) Collaborative Study Group at New York City/Bronx Consortium which was funded by the National Institute of Allergy and Infectious Diseases UO1-AI-35004. We are indebted to the participants of this study, many of whom have now devoted over 15 years of their life to this effect. We also thank Howard Strickler, Xiaonan Xue, Steve Gange, Peter Bacchetti, Minge Xie & Zhiqiang Tan for their helpful comments.

## ABBREVIATIONS

AR(1): –Autoregressive Order 1
BS: – Between Subject
BUN: – Blood Urine Nitrogen
Co-DOSE: – <u>Co</u>nditionally <u>D</u>ependent <u>O</u>n <u>S</u>ibling <u>E</u>xposure
CS: – Cross Sectional
E: – Equicorrelation
EGFR: – Estimated Glomerular Filtration Rate
GEE: – Generalize Estimation Equations
IND: – Independent
MA(2): – Moving Average Order 2
MM: – Mixed Models
WIHS: – Women’s Interagency HIV Study
WS: – Within Subject

## APPENDIX 1– Homology Between Co-DOSE (4) Occurrence and Between and Within Subject Slopes being the Same or Differing

Figure 5 illustrates using the example of Section 2B (with *K* = 1) that Co-DOSE (4)” occurs if *β_BS_* ≠ *β_WS_*. Remember that in this example, *β_BS_*=0.9, *β_WS_*=3, *β_CS_*=1.60. Now let *J*=2. So for the Between / Within Subject decomposition model; 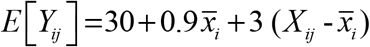. If the overall mean of *X_ij_* for all repeat measures in the sample was 180 (i.e. 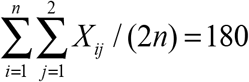) then the full cross sectional model is *E*[*Y_ij_*] = −96+1.60(*X_ij_*). If a subject’s two weight measures are *X_i1_* and *X_i2_* =220, then for the first measure, the Cross Sectional model estimates *E*[*Y_ij_*|*X*_*i*1_] = −96 + 1.60 (200) = 224. But since *X_i2_* =220 and 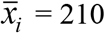, as we saw earlier Between/Within Subject decomposition gives; *E*[*Y_ij_* | *X*_*i*1_,*X*_*i*2_] =30+0.9(210)+3(200-210) = 189. Thus *E*[*Y_1j_*|*X_i1_*] is not independent of *X_i2_* since X_i2_ is informative of where 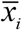 falls and the slope for 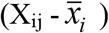 is different than the slope for 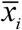 when *β_BS_* ≠ *β_ws_*. But if *β_BS_* = *β_WS_*, *X_i2_* is non-informative on *Y_i1_*|*X_i1_* as E[*Y_ij_*] = α_CS_ + β_CS_ 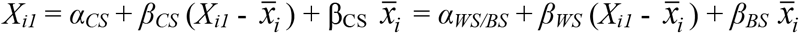 since *β_BS_ = β_WS_*.

**Figure 5.**
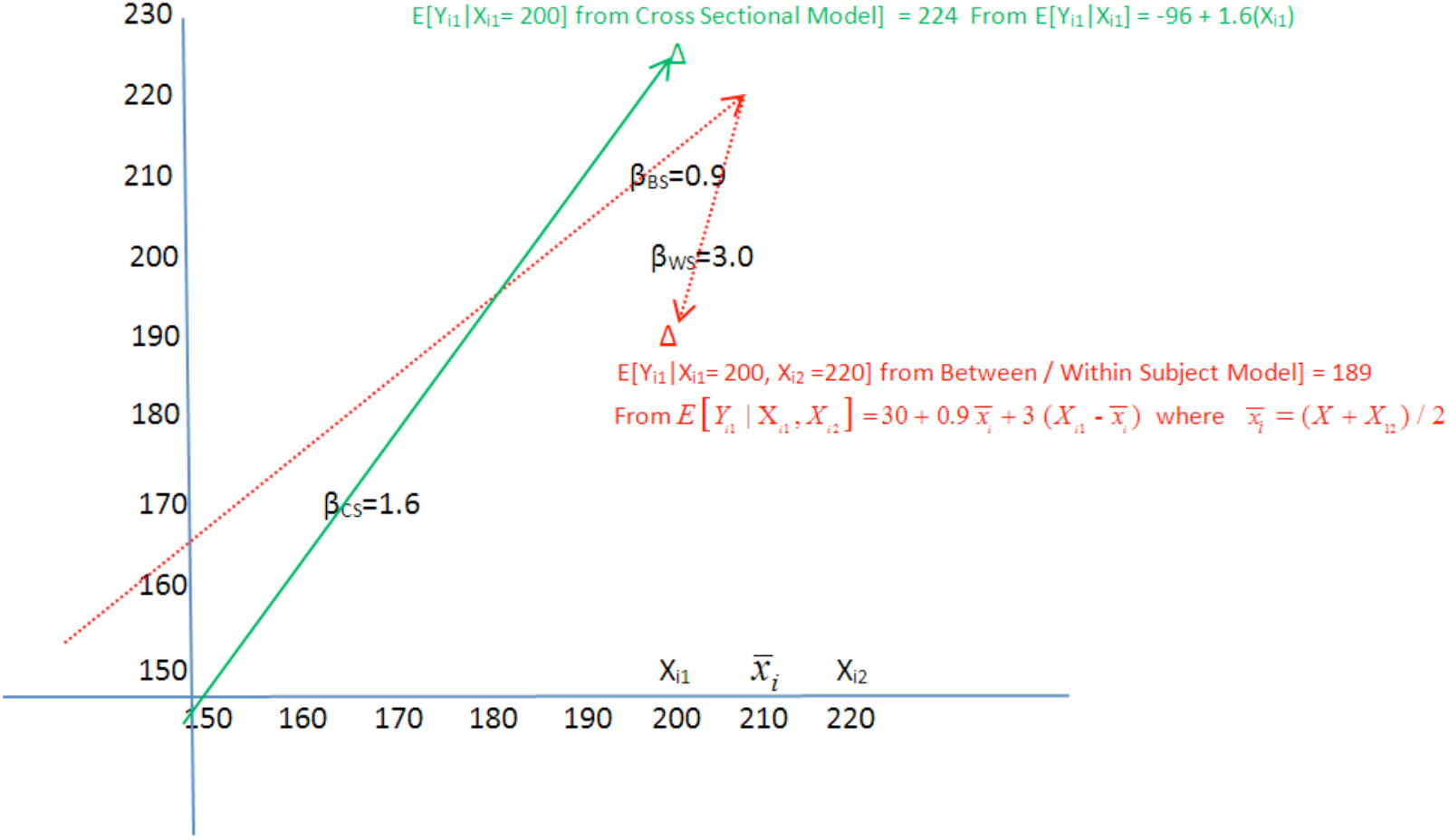
Illustration that E [Y_i1_}X_i1_] is not independent of X12 in Cross Sectional Regression for K=1 when B_BS_ ≠ β_WS_ in a Between/Within Decomposition Model with J_i_ = 2, X_i1_=200 and X_i1_2=220

As J_i_=2, in the prior example, the second observation was deterministic for 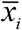. But for J_i_ > 2, additional *X_ij’_* go into computation of 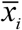 are thus still informative on relative contributions of 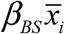 and 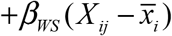 on E[*Y_ij_*|*X_ij_*]. For β_WS_ not well defined, additional *X_ij_’* may still be informative on a 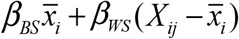 decomposition with *β_WS_* = *E*[*β_WS_*].

Whether or not Co-DOSE (4) occurs also informs if *β_BS_* = *β_WS_*. If for a given *j, E*[*Y_ij_* | *X_ij_*]is independent of all other *X_ij’_*, then *E*[*Y_ij_* | *X_ij_*]is independent of 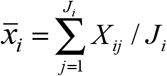, which only happens if *β_BS_* = *β_WS_*. But if *E*[*Y_ij_* | *X_ij_*]is not independent of other *X_ij’_*, then; i) if 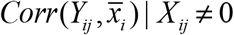, then *β_WS_* (if well defined) ≠ *β_BS_*, ii) otherwise if 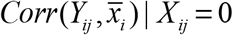, then *β_Ws_* is not well defined.

## APPENDIX 2– Illustration that Measurement Error Which is Independent of the Outcome Pushes *β_WS_* and *β_BS_* to Zero With Greater Impact on *β_WS_*

To illustrate this for *K*=1, let there always be the same number of replicates, *J*, per subject and assume in the absence of Measure Error, *β_WS_* = *β_BS_* = *β_CSR_*. For example, let 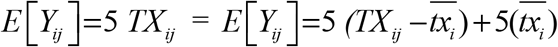, for simplicity the intercept is 0. But we observe *X_ij_* = *TX_ij_* + *M_ij_* where *M_ij_* is Measure Error with variance 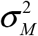 that is independent across all *i*’s and *j*’s and also from *Y_ij_*. Let *TX_ij_* vary with *j* within *i* as follows; *TX_ij_* = *TC_i_* + *TR_ij_* where *TC_i_* is a central tendency of *TX* for subject *i*, while *TR_ij_* is within subject *i* repeated visit variation in *TX* across the *j*’s. Let 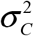 and 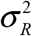 be variances of *TC_i_* and *TR_ij_*, respectively.

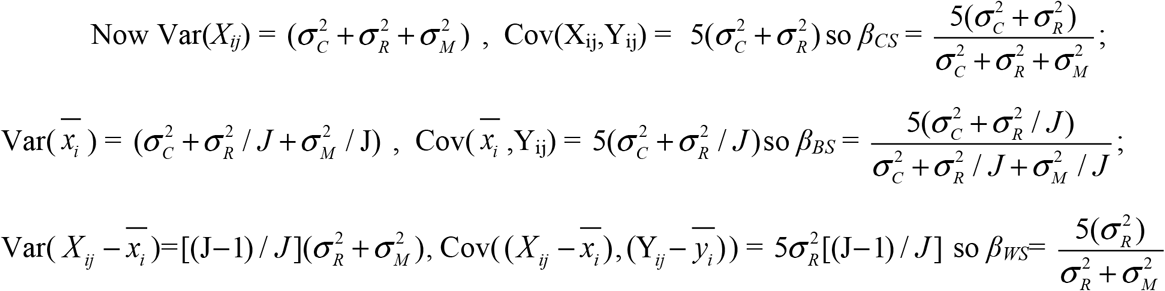

For example, let 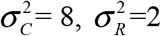 and 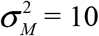 and *J*=5. The entire variance of *X_ij_* is 20 of which half, 10, is from Measure Error, 8 is variation of the central tendency of *X* between subjects and 2 is variation of *X* within subject. Then *β_CS_* = 5(10)/20 = 2.5, *β_BS_* = 5(8.4)/10.4 = 4.03 and *β_WS_* = 5(2)/12 = 0.83. Considering that without Measure Error, true Between and Within person slopes both = 5, Measure Error has greatly tempered *β_WS_*=0.83 towards 0 while *β_BS_*=4.03 is the least tempered. This happens because *β_BS_* most fully retains the signal in *X*, but tempers *M* through averaging, while *β_WS_* more fully retains *M* while excluding the between subject signal in *X*.

